# Identification of a unique ZIP transporter involved in zinc uptake via the arbuscular mycorrhizal fungal pathway

**DOI:** 10.1101/2020.09.28.317669

**Authors:** Stephanie J Watts-Williams, Stefanie Wege, Sunita A Ramesh, Oliver Berkowitz, Matthew Gilliham, James Whelan, Stephen D Tyerman

## Abstract

Low soil zinc (Zn) availability is a limiting factor for crop yield, and increasing Zn content is a major target for the biofortification of major crops. Arbuscular mycorrhizal (AM) fungi associate with the roots of most terrestrial plant species and improve the host plant’s growth and nutrition through the mycorrhizal pathway of nutrient uptake. Although the physiology of Zn uptake through the mycorrhizal pathway is well established, the identity of the molecular components responsible for Zn transport in the mycorrhizal pathway are unknown.

RNA-seq analysis identified the putative Zn transporter gene *MtZIP14* by its marked up-regulation in *Medicago truncatula* roots when colonised by the AM fungus *Rhizophagus irregularis* under varying soil Zn supply. Expression of GFP-tagged MtZIP14 in roots revealed that it is exclusively localised to the site of plant-fungal nutrient exchange in cortical cells, the peri-arbuscular membrane. Expression of MtZIP14 in a yeast mutant lacking Zn transport function restored growth under low Zn availability. *M. truncatula MtZIP14* loss-of-function mutants had reduced shoot biomass compared to the wild-type when colonised by AM fungi and grown under low Zn. Vesicular and arbuscular colonisation, but not hyphal colonisation, were also lower in *mtzip14* mutant plants.

Based on these results we propose that MtZIP14 plays a key role in the transport of Zn from AM fungus to plant across the peri-arbuscular membrane, and *MtZIP14* function is crucial to plant competitiveness in a low Zn soil.

**Significance statement:** Majority of crop plant species associate with arbuscular mycorrhizal fungi, which can increase plant nutrient uptake. Improving our knowledge of how Zn is taken up in mycorrhizal plants will lead to improved plant and human Zn nutrition outcomes. Here, we report a novel plant transporter with a major role in Zn nutrition of mycorrhizal plants. MtZIP14 is involved in Zn transport, is exclusively localised to the specialised plant-fungal interface in roots, and impairment of *MtZIP14* gene function results in negative impacts on both plant growth and Zn nutrition.

## Introduction

Zinc (Zn) is an essential co-factor for >300 enzymes in plants, making it critical for processes such as carbon fixation, transcription, and production of ATP (1, 2). It is also an essential micronutrient for humans and Zn deficiency is the fifth leading risk factor for disease in developing countries with high mortality (3). Zinc is taken up at the plant-soil interface in its divalent form Zn^2+^ by the Zn-regulated iron-regulated transporter-like protein (ZIP) family, which also have a role in the transport of other transition metals (4). ZIP transporters are involved in cellular Zn homeostasis (5, 6) and the plant response to Zn deficiency (7, 8), while over-expression of ZIPs can lead to increased tissue Zn concentrations (9, 10). Characteristics of most ZIP transporters include eight predicted transmembrane-spanning α-helices, and a hydrophilic variable region between helix III and IV that contains a potential metal-binding domain (4). In the model legume *M. truncatula*, 16 predicted ZIP transporters have been identified through phylogenetic analysis (11) and four of those have been characterised for Zn transport function by expression in the yeast mutant ZHY3 that lacks Zn transporters (12); however, the specific roles of these ZIPs *in planta* is currently unknown.

The majority of terrestrial plant species form associations with arbuscular mycorrhizal (AM) fungi; resource exchange is critical to the symbiotic association and typically involves trade of inorganic nutrients from the fungus and carbon resources from the plant (13). A primary benefit of colonisation by AM fungi is an improvement in plant growth and nutrition, particularly of nitrogen (N), phosphorus (P) and Zn nutrition (14, 15). Managed effectively, AM fungi provide a tool for improved crop Zn nutrition in the field, particularly on Zn-deficient soils (16). Radioisotope tracing studies have demonstrated that the AM fungus *Rhizophagus irregularis* can contribute as much as 25% of shoot Zn uptake in tomato plants, 24% of grain Zn in wheat, and 12% in barley (17, 18).

Considerable progress has been made toward identifying the components involved in P and N AM associations (19-21), and a AM-specific plant Cu transporter has been recently identified (22). In order to fully exploit the AM symbiosis for improved agricultural outcomes (i.e. crop quantity and quality, biofortification), it is essential that these molecular components are identified (23, 24). While an AM fungal transporter that facilitates Zn regulation in extraradical hyphae has been identified (25), no plant Zn transporter has been identified that is involved in the AM association (23, 26). It has been postulated that an, as yet, unidentified Zn transporter is exclusively located on the plant-derived peri-arbuscular membrane (PAM) present in AM-colonised root cortical cells, responsible for the import of Zn^2+^ supplied by the fungus (19). Here, we propose that MtZIP14 is this postulated transporter.

We discovered MtZIP14 through a transcriptomic screen and through functional characterisation identified it as a novel Zn transport protein exclusively expressed in roots upon AM colonisation; and is localised to the PAM. Zn transport capacity of MtZIP14 was supported by heterologous expression in yeast. Examination of *M. truncatula* loss-of-function mutants demonstrated a negative effect on shoot biomass and AM colonisation of roots, which was linked to the transport of Zn. We have presented evidence that MtZIP14 has a critical role in Zn transport by the AM pathway of uptake into plants

## Results

### Identification of MtZIP14 as a candidate Zn transporter specifically expressed in AM colonised roots

To identify genes potentially involved in the mycorrhizal uptake of Zn we performed a RNA-seq experiment using *Medicago truncatula* grown at different soil Zn concentrations and inoculated with the AM fungus *Rhizophagus irregularis* compared to mock inoculation (Figure S1a; Table S1). In order of increasing soil Zn addition (0, 5, 20 mg Zn kg^-1^), there were 589, 201 and 918 genes that increased significantly in abundance with AM colonisation (Figure 1a) and 221, 33 and 159 genes that decreased in abundance (Figure 1b). At 0 mg kg^-1^ added Zn, the transcripts up-regulated by AM colonisation were associated with the GO terms copper, iron, and manganese ion binding, confirming the efficiency of the AM inoculation and low nutrient soil. As expected, non-colonised plants displayed marked changes in transcript abundance with Zn-deficiency (Zn 0) (135 up, 538 down) in comparison to the AM-colonised plants (29 up, 37 down).

**Figure 1.**
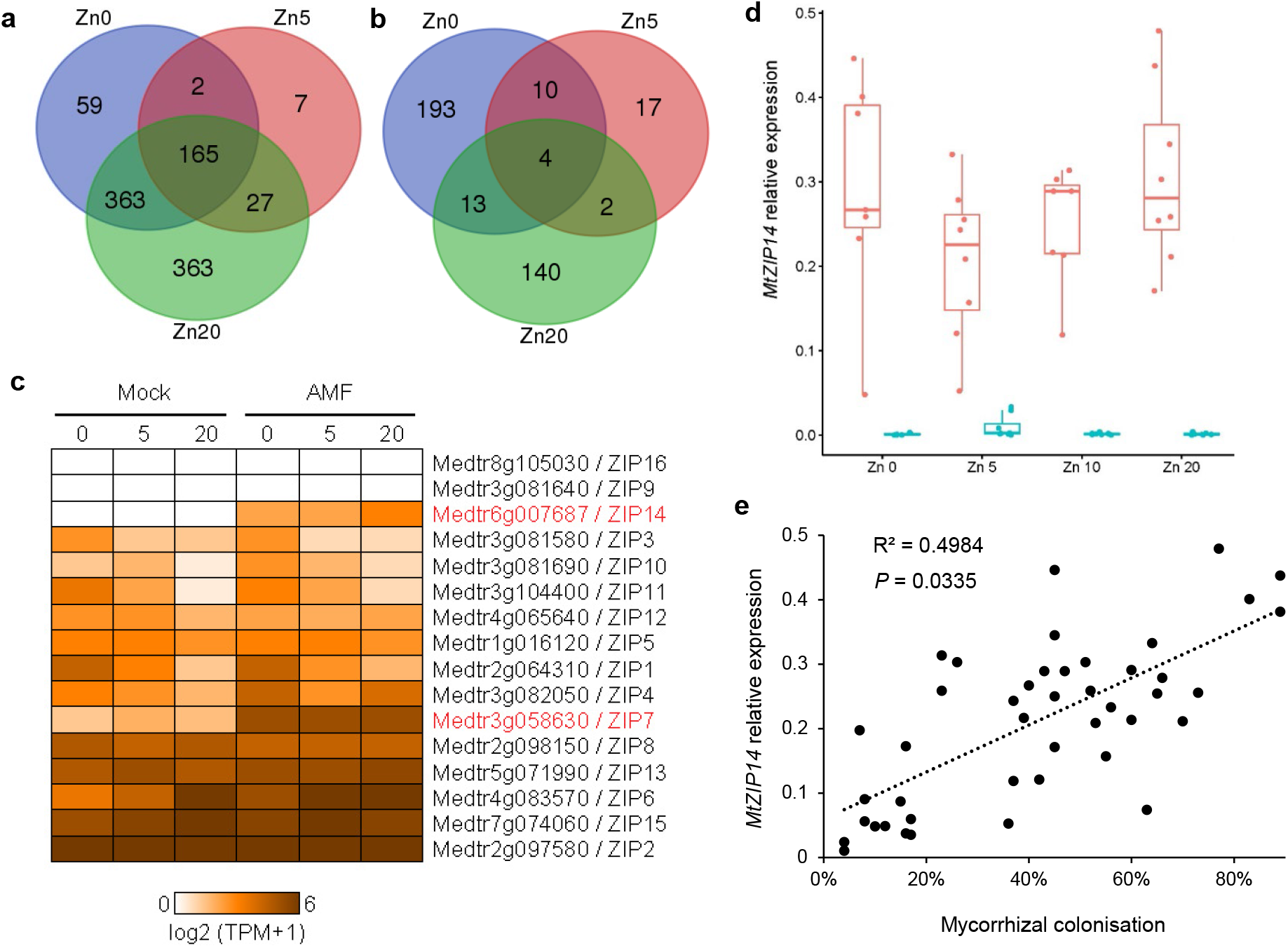
Numbers of significantly up-regulated (a) and down-regulated (b) *Medicago truncatula* A17 genes by *Rhizophagus irregularis* colonisation, expression of 16 genes annotated as ZIP transporters (c) split into three soil Zn addition treatments: Zn0 no addition; Zn5 5 mg kg^-1^ addition; Zn20 20 mg kg^-1^ addition. Genes highly up-regulated in AM colonised plants across all three Zn addition treatments are highlighted in red. Expression of *MtZIP14* in the roots of *Rhizophagus irregularis* -inoculated (pink) and mock-inoculated (blue) plants grown at four soil Zn additions (d) and relationship between root length colonised by AM fungal structures assessed by microscopy and *MtZIP14* gene expression (e). Gene expression is calculated as the gene-of-interest relative to the geometric mean of three housekeeping genes (Table S5).

Three lists of candidate genes with a potential role in AM fungal Zn nutrition were compiled based on their gene annotation as zinc transporter (ZIP), heavy metal transporter or zinc-binding (Table S2-4). Of all 16 annotated *ZIP* genes in *M. truncatula*, only one (*MtZIP14; Medtr6g007687*) was exclusively expressed in AM colonised root cells independent of the Zn concentration, and another gene was up-regulated in all Zn treatments (*MtZIP7; Medtr3g058630*), while all others showed no AM specificity (Figure 1c). A previous study had shown that the MtZIP7 is a manganese (Mn) transporter (12), suggesting that MtZIP7 is not primarily involved in Zn uptake. The only remaining and most promising candidate, *MtZIP14*, was uncharacterised, and showed an expression pattern consistent with involvement in the AM pathway of Zn uptake. Quantitative RT-PCR on samples from an independent experiment confirmed that *MtZIP14* is almost exclusively expressed in plants that have been colonised by *R. irregularis* (Figure 1d). Expression of *MtZIP14* was not affected by increasing soil Zn concentration (0, 5, 10, 20 mg kg^-1^ added Zn), and there was a positive (R^2^ = 0.498), significant (*P* = 0.03) relationship between the root colonisation by AM fungi and expression of *MtZIP14*, suggesting that increased colonisation by AM fungi is associated with increased expression of *MtZIP14* (Figure 1e).

Two genes encoding HMA-domain proteins contained in the heavy metal transporter list were up-regulated by AM colonisation: one in Zn 0 and 5 (*Medtr0041s0140*) and one in all Zn treatments (*Medtr6g051680*) (Figure S1b). HMA-domain proteins play key roles in transporting monovalent and divalent ions in plants, and in detoxification (27). In the zinc-binding candidate list there was a Zn-binding dehydrogenase oxidoreductase gene up-regulated in all Zn treatments (*Medtr8g035880*) (Figure S1c); Zn-binding alcohol dehydrogenases catalyse the reduction of acetaldehyde to ethanol, mainly in meristematic tissues such as root apices under anaerobic conditions (2). The expression of these three genes were determined in an independent experiment using quantitative RT-PCR; the HMA-domain protein *Medtr6g051680* was induced by AM colonisation across all Zn conditions (Figure S2a), while *Medtr0041s0140* was down-regulated by AM colonisation in this experiment (Figure S2b). The Zn-binding dehydrogenase oxidoreductase gene (*Medtr8g035880*) was exclusively expressed in AM colonised roots in all the soil Zn treatments (Figure S2c).

### Characterisation of MtZIP14

We concentrated on further characterising *MtZIP14*, as it was the most likely candidate of the AM up-regulated genes to transport Zn across the PAM. *In silico* analysis predicted a potential metal-binding domain rich in histidine residues between transmembrane III and IV, similar to the other ZIP proteins with Zn-transport function characterised in *M. truncatula* (MtZIP1, 2, 5, 6; (12, 28)) (see protein sequence alignment Figure S3).

Plant transporters involved in export or import of nutrients and compounds traded between the fungus and the plant are typically exclusively localised to the specialised PAM. We therefore investigated the subcellular localisation of MtZIP14 through a C-terminal translational fusion with GFP introduced into *Agrobacterium rhizogenes* ARqua1. The GFP-tagged construct was introduced into *M. truncatula* A17 through hairy roots transformation (29); transformed plants were subsequently inoculated with the AM fungus *R. irregularis*. Confocal imaging revealed GFP fluorescence exclusively localised to the fine branches of arbuscules in the cortical cells of roots (Figure 2a). This fluorescence pattern suggested that MtZIP14 expression is specific to colonised cells, and subcellular localisation is specific to the PAM section of the plant cell plasma membrane. No GFP fluorescence was detected in cells not colonised with *Rhizophagus irregularis* or in the arbuscules from plants transformed with the empty vector control (Figure 2b), strongly supporting the conclusion that MtZIP14-GFP is PAM-localised.

**Figure 2.**
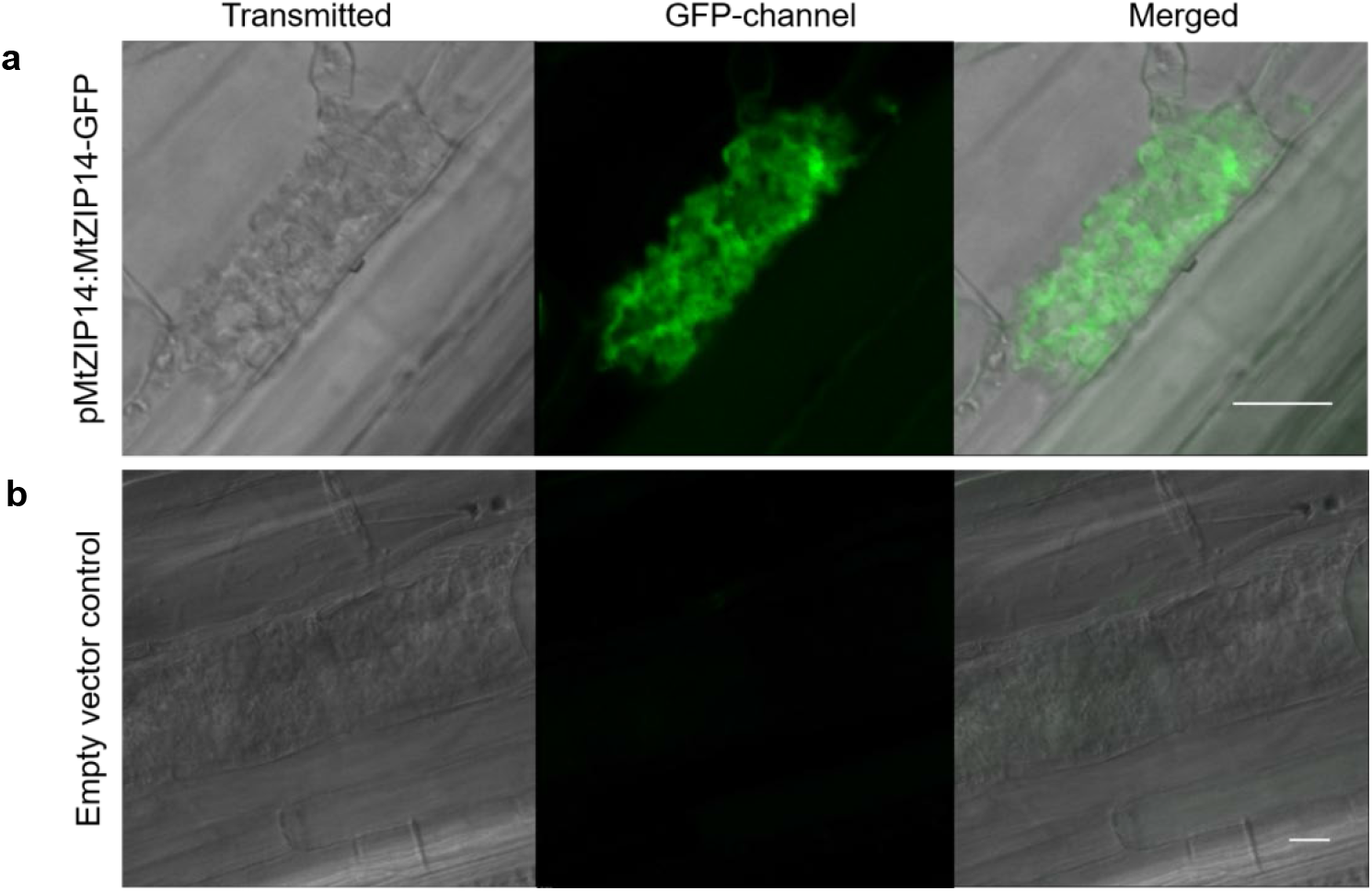
Confocal laser-scanning microscopy images of C-terminally GFP-tagged MtZIP14, with fluorescence shown in green (a), and the empty vector control (b) in cortical cells of *Medicago truncatula* A17 containing arbuscules formed by *Rhizophagus irregularis* colonisation. Scale bar is 10 µm.

To test for the involvement of MtZIP14 in Zn transport, *MtZIP14* was cloned into a heterologous expression system – yeast (*Saccharomyces cerevisiae*) lacking Zn transporters; the yeast strain ZHY3 (*zrt1zrt2;* (30, 31)) displays reduced growth under low Zn conditions. In the EDTA-only YNB media, neither the empty vector control nor the MtZIP14 expressing ZHY3 yeast strain grew, confirming the growth defect of ZHY3 (*zrt1zrt2*) mutant (Figure 3a). Interestingly, already in the lowest Zn addition concentration (0.1 mM) where Zn^2+^ was available at nanomolar concentration, the MtZIP14-expressing yeast grew, while the empty vector ZHY3 strain did not (Figure 3b), suggesting that MtZIP14 is able to mediate Zn uptake from very low external Zn concentrations and is likely a high affinity Zn transporter. In the 0.2 and 0.5 mM added Zn EDTA-YNB (Figure 3c,d), the empty vector displayed slow growth with OD_600_ increasing after 35 hours, suggesting that higher Zn is sufficient to enable this yeast strain to survive. Expression of MtZIP14 significantly increased yeast growth over the empty vector control, which was especially evident at 0.5 mM Zn, where MtZIP14 growth peaked (Figure 3g). However, growth of the MtZIP14-expressing yeast was reduced at 1 mM added Zn compared to the empty vector control, suggesting the transport of Zn via MtZIP14 resulted in Zn influx to toxic concentrations and inhibited growth of the yeast (Figure 3e). At the highest Zn addition (1.5 mM), the empty vector yeast grew well but the MtZIP14-expressing yeast did not grow until 35 hours, and growth thereafter was poor, evidencing further the toxicity hypothesis, and that MtZIP14 may be a dual-affinity transporter (Figure 3f). Growth of ZHY3 on the solid YNB agar media for 96 hours followed the same pattern as the liquid YNB; MtZP14-expressing yeast grew in all Zn treatments and best at the 0.5 mM added Zn (Figure S4a-c), while the empty vector yeast grew well only at 1mM added Zn. The wild-type (WT) positive control yeast strain (DY1457) grew on all solid agar experimental conditions with Zn addition (Figure S4a-c). This data suggest that MtZIP14 is a membrane protein and able to facilitate Zn transport into cells.

**Figure 3.**
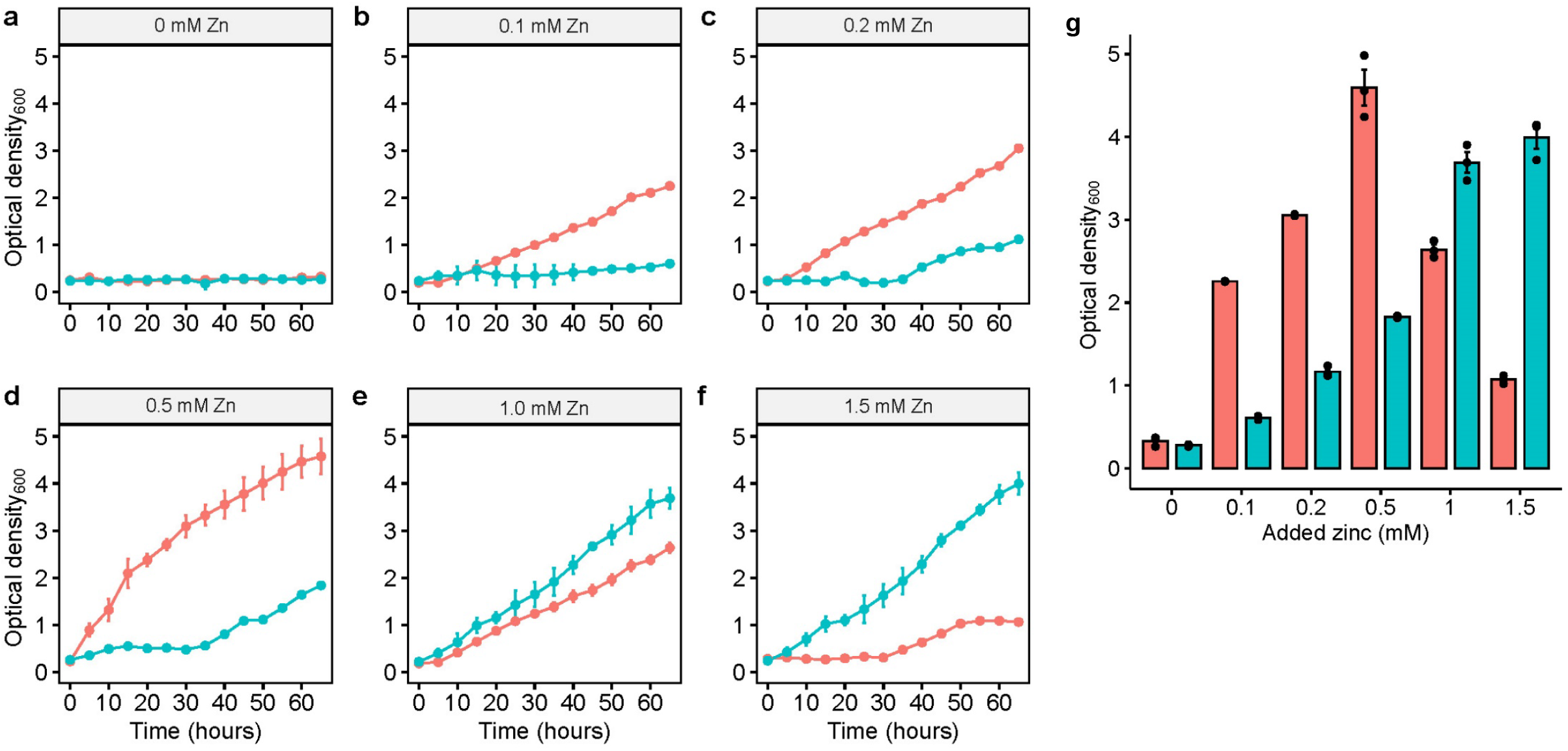
Complementation of the Zn transporter (*zrt1zrt2*) mutant (ZHY3) yeast strain with MtZIP14 (pink), or with the empty vector pDEST52 (blue), grown over a 66 h period in liquid YNB -uracil media with 2% galactose and 1 mM EDTA. With the addition of EDTA only (a), there was no growth of any yeast strains without Zn supplementation. The Zn supplementation treatments were 0.1 (b), 0.2 (c), 0.5 (d), 1.0 (e) and 1.5 (f) mM ZnSO_4_. After 66 h of growth the final OD_600_ all treatments was recorded (g). Values are mean ± standard deviation of the mean, *n*=3.

*M. truncatula* plants with Tnt1 retrotransposon insertion in the *MtZIP14* gene were isolated and analysed. The generated *mtzip14* plants had either no detectable expression of *MtZIP14* (NF8057; knock-out, KO) or a strongly reduced expression (NF4665; knock-down, KD) to approximately one third of the out-segregated WT (Figure 5a). As a control, we used out-segregated WT plants from those two lines, which expressed *MtZIP14* when colonised by the AM fungus *R. irregularis* while the mock-inoculated plants had no expression, confirming the results obtained with WT plants in the RNA-seq experiment.

**Figure 4.**
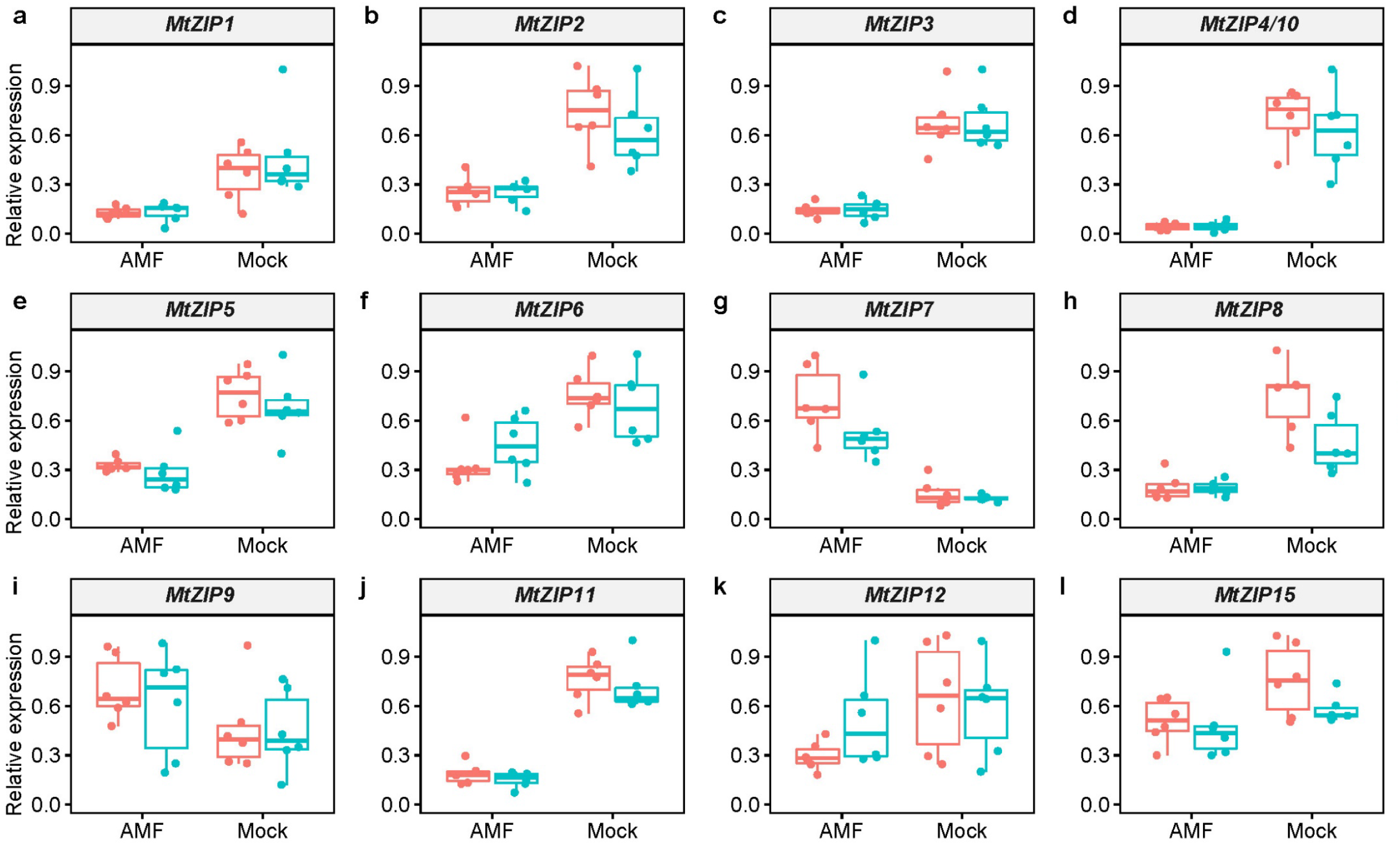
Expression of 12 *ZIP* transporter genes (a-j), in the roots in the *mtzip14* (pink) and segregating WT (blue) plants grown with or without inoculation by the AM fungus *Rhizophagus irregularis*. Gene expression is calculated as the gene-of-interest relative to the geometric mean of two housekeeping genes (Table S5).

**Figure 5.**
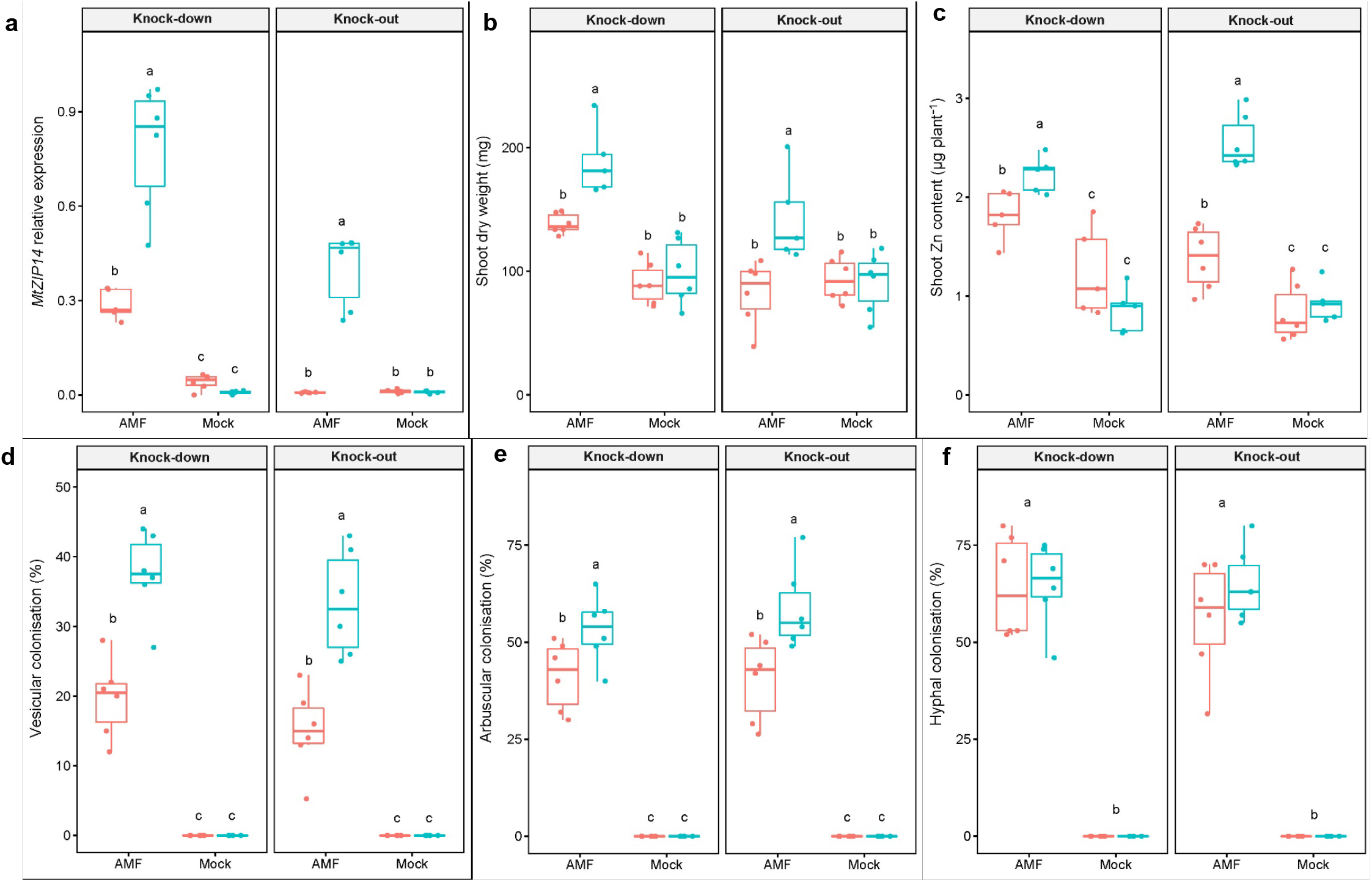
Root expression of *MtZIP14* (a), shoot dry weight (mg) (b), shoot Zn content (µg Zn plant^-1^) (c), root length colonised by AM fungus *Rhizophagus irregularis* in terms of percentage vesicles (d), arbuscules (e) and internal hyphae (f) in the knock-out and knock-down *mtzip14* (pink) and segregating WT (blue) lines grown with or without inoculation by the AM fungus *R. irregularis*. Means with different letters are considered significantly different (*P*<0.05) as per Tukey’s HSD *post hoc* test. Where one letter appears above two boxes, it represents a significant main effect of *Mycorrhiza* where the two genotypes are pooled. Gene expression is calculated as the gene-of-interest relative to the geometric mean of two housekeeping genes (Table S5).

We first investigated how the loss of *MtZIP14* function altered the expression of other *ZIP* transporter genes by analysing 15 additional *MtZIP*s by quantitative RT-PCR from the KO genotype (NF8057) roots and the corresponding WT. Majority of the *ZIP* genes measured were highly down-regulated in the AM colonised roots compared to the mock-inoculated roots, regardless of genotype (i.e. both *mtzip14* and WT were similarly down-regulated in AM colonised roots) (Figure 4a-l; Table S6). As expected, *MtZIP7* was the only *ZIP* gene found to be up-regulated by AM colonisation, and we found that it is more highly up-regulated in *mtzip14*, suggesting a transcriptional impact on *MtZIP7* due to the loss of *MtZIP14* (Figure 4g). No transcripts were detected for *MtZIP13* or *ZIP16*.

We then examined the loss of *MtZIP14* function on the plant and AM fungal phenotypes. The shoot biomass of both *mtzip14* plant lines was reduced when compared to the out-segregated WT plants when colonised by *R. irregularis* under Zn deficient conditions (Figure 5b; Table S7; Table S8), with no significant difference for the mock-inoculated plants. Shoot Zn concentrations (mg kg^-1^) were increased by AM inoculation in all plants (Figure S5); meanwhile, colonised *mtzip14* plants had lower Zn content (µg Zn per plant) than the WT plants, whereas, the mock-inoculated *mtzip14* and WT plants contained similar amounts of Zn (Figure 5c).

For both *mtzip14* lines, vesicular (Figure 5d) and arbuscular (Figure 5e) colonisation were both lower than the WT, while hyphal colonisation of roots was not significantly different (Figure 5f). There was no colonisation by AM fungi in the mock-inoculated plants; and shoot biomass, Zn nutrition and root AM colonisation were comparable in the R108 wild-type plants and segregated WT lines (Figure S6a-f).

We then conducted a principal components analysis (PCA) to analyse all data simultaneously. PCA revealed a marked effect of *Mycorrhiza* treatment on the plants when all plant physiological response variables were considered together (Figure S7). However, when the AMF and Mock data were analysed separately, there was a significant separation based on *Genotype* in the AM colonised plants only (Figure 6a,b). Furthermore, the loadings show that WT plants were separated from the *mtzip14* plants by their greater AM colonisation (arbuscular, vesicular, and hyphal) and shoot Zn contents. Shoot Zn contents were also highly correlated with vesicular colonisation of the roots, and not to the contents of other nutrients. Taken together, the PCA provides evidence of the link between AM fungi and plant Zn nutrition in the context of *MtZIP14* function.

**Figure 6.**
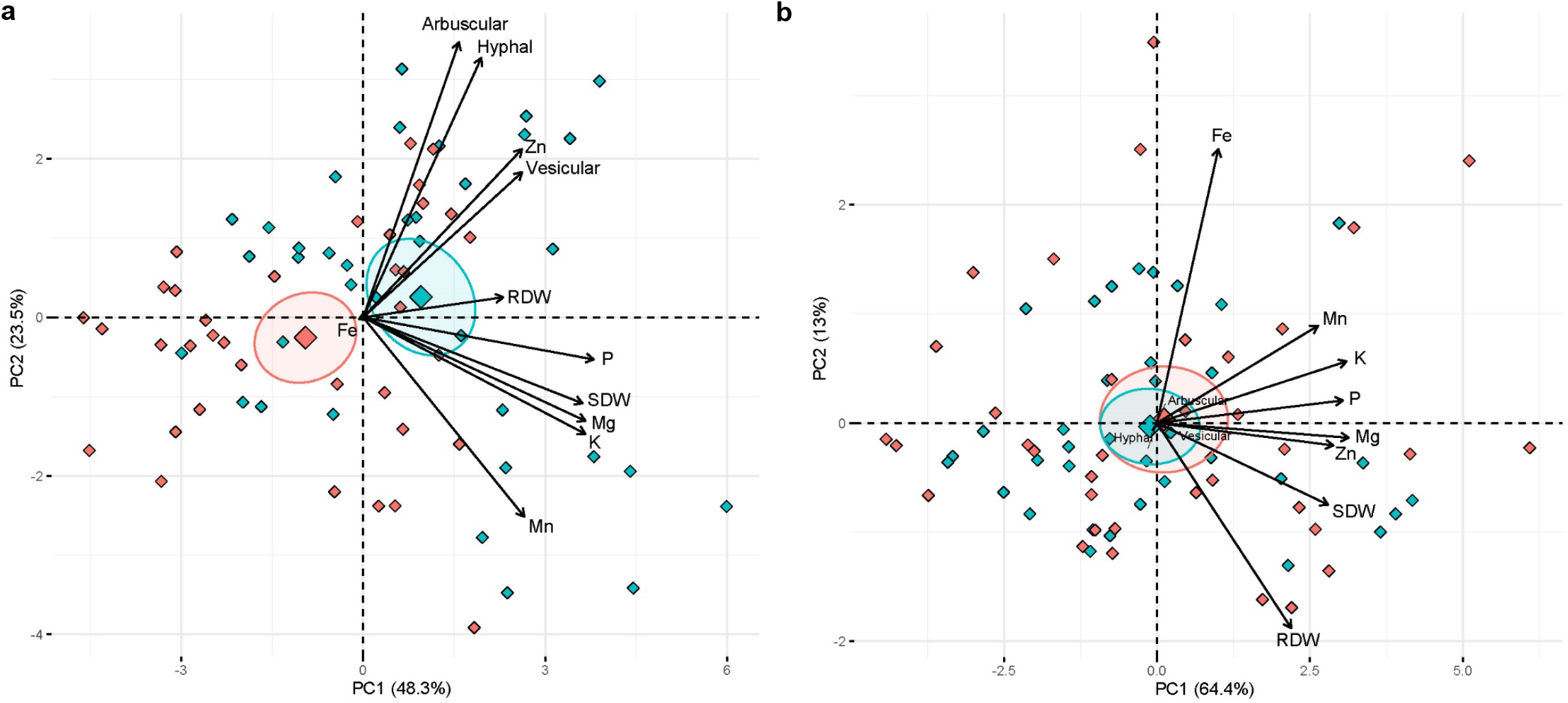
Principal components analysis (PCA) biplot displaying scores in the first two principal components (PC1: x-axis, PC2: y-axis) following PCA of biomass (shoot and root dry weights), shoot nutrient contents (µg plant^-1^; Zn, P, Mg, K, Mn, Fe) and arbuscular mycorrhizal colonisation (arbuscular, vesicular, hyphal) response variables of *mtzip14* mutant plants (pink) and segregating wild-type plants (blue) grown in three replicated experiments. Plants were either inoculated with the arbuscular mycorrhizal fungus *Rhizophagus irregularis* (a) or mock-inoculated (b). The sign and magnitude of the contribution of variables is indicated by the loadings (arrows). The large diamonds signify the mean and 95 % confidence ellipse for each *Genotype* treatment.

## Discussion

### A role for MtZIP14 in arbuscular mycorrhizal Zn uptake

We have identified MtZIP14 as a plant transport protein with a major role in the uptake of Zn via the AM pathway of uptake. When grown in a Zn-deficient soil, the WT with functional *MtZIP14* had a clear benefit over the *mtzip14* KO and KD lines, and WT shoots produced significantly more biomass. This indicates that the function of *MtZIP14* is critical to the colonised plant being competitive in a Zn-deficient soil. Furthermore, the PCA indicates that of all measured nutrients, shoot Zn contents were affected the most by the disruption of *MtZIP14*, demonstrating a role for *MtZIP14* in mycorrhizal plant Zn uptake. It also suggests that the non-colonised control plants could not compensate the loss of AM-derived Zn with increased uptake via the direct pathway (i.e. root uptake from the rhizosphere) to reach similar Zn contents as the AM-colonised WT plants. Non-colonised plants took up only around ∼50 % of the Zn compared to the colonised WT plants. Gene expression analysis of the KO and KD lines revealed that the mock-inoculated plants showed higher expression of at least eight *ZIP* genes compared to the AM colonised plants, including four ZIP transporters that have been shown to transport Zn in yeast previously (MtZIP1, 2, 5, 6); this suggests that expression of non-PAM *ZIP* genes are generally suppressed in AM plants. Despite the general down-regulation of *ZIP* transporter genes in the AM-inoculated plants, the Zn concentrations of the AM inoculated plants were still greater. This might suggest that Zn uptake via the AM pathway is strongly preferable to the plant, compared to direct uptake.

AM colonised *mtzip14* plants accumulated more Zn in their shoots compared to the non-colonised *mtzip14* plants. This indicates that the *mtzip14* plants still had a Zn uptake advantage by being colonised by AM fungi, although not to the extent of the WT plants with a functional *MtZIP14*. The source of the advantage may be another AM-specific transporter, besides MtZIP14, that is able to transport Zn across the peri-arbuscular membrane, which is also expressed in non-colonised plants and was therefore not identified in our RNA-seq. In addition, *MtZIP7* was expressed more highly in the *mtzip14* plants than the WT plants, which may suggest that MtZIP7 might be able to compensate partially for the loss of *MtZIP14* function. MtZIP7 may be able to also transport Zn at a low affinity or low rate, and increased expression and protein abundance might therefore lead to increased Zn uptake in *mtzip14*. Alternatively, the effect may be due to indirect environmental effects of the AM symbiosis on the availability of Zn in the soil (e.g. through exudation that mobilises Zn in soil) that led to increased plant uptake of Zn via the direct pathway.

Loss of *MtZIP14* function affected the colonisation of the roots by *R. irregularis* suggesting an important role for the transporter in plant-fungus communication. The proportion of ‘functional’ AM structures (arbuscules and vesicles) were lower in the *mtzip14* mutant plants, while hyphal colonisation was not significantly different, indicating that MtZIP14 is important for the correct formation of fungal structure within the root, but not the colonisation event itself. A similar phenotype was observed in plants lacking the AM-specific Pi transporter gene (*MtPT4*) function (20), and rice plants lacking a symbiotic nitrate transporter gene (*OsNPF4*.*5*) function (21). This suggests that the plant-fungal symbiosis is somewhat disrupted by the loss of *MtZIP14* expression, and that the active sites of nutrient transport (arbuscules), as well as fungal resource storage units (vesicles), were not able to be produced by the fungus to the same extent due to this disruption.

We observed that expression of *MtZIP14* was not down-regulated in AM-colonised plants when Zn was in high supply, which correlates with Zn isotope data in tomato showing that the mycorrhizal pathway of Zn uptake is not supressed at high Zn concentrations, and is similar regardless of available Zn in the soil (18). This is in contrast to the expression of the AM-specific Pi transporter *MtPT4*, which is strongly down-regulated when P is highly available to the plant (19), and the transport of isotope labelled P via the mycorrhizal pathway of uptake is likewise suppressed (32). Divalent cation transporters such as ZIPs are often less selective compared to other nutrient transporters, and uptake via the AM pathway might be beneficial to the plant, as compared to the direct pathway, which could include the risk of importing unwanted cations such as cadmium (Cd). This may also explain the general down-regulation of non-PAM ZIPs in AM-colonised plants.

We have identified and described a ZIP transporter that has an important role in Zn transport into plants via the mycorrhizal pathway. This contributes to the development of a comprehensive plant-AM fungal nutrient exchange model across the PAM, and will stimulate research into identifying other micronutrient transporters involved in the AM pathway of nutrient uptake.

## Methods

### Plant growth conditions and harvest

The *M. truncatula* plants grown for RNA-sequencing, gene expression, protein localisation and loss-of-function phenotyping were all grown in similar conditions; briefly, seeds of *Medicago truncatula* ecotypes A17 or R108 (loss-of-function studies only) were surface-sterilised, surface-scarified lightly with sandpaper, imbibed and germinated on filter paper as previously described (33).

Pre-germinated seedlings were moved into pots inoculated with the AM fungus *Rhizophagus irregularis* WFVAM10, or mock-inoculated. The growth substrate was a mix of autoclaved fine sand mixed in a ratio of 9:1 with sieved and autoclaved low nutrient soil from the Mallala region of South Australia. The final soil/sand mix had a plant-available (DTPA-extractable) Zn concentration of 0.3 mg Zn kg^-1^. The *R. irregularis* inoculum comprised dry soil, root pieces, spores and hyphae from a pot culture where *R. irregularis* was previously cultured on Marigold (*Tagetes patula*) seedlings for 12 weeks. The mock inoculum was cultured in exactly the same way but without the addition of *R. irregularis* to the culture. For each pot, 630 g of the sand/soil growth substrate was mixed with 70 g of the AM fungal or mock inoculant substrate prior to transplantation. Plants were grown in a controlled environment chamber with day/night conditions set at 24 °C/20 °C and 16/8 hours of light/dark. Plants were watered until draining with reverse osmosis (RO) water three times per week. In order to ensure the only limiting plant essential nutrient was Zn, plants were given 10 mL each of a modified Long-Ashton solution with Zn omitted from the micronutrient cocktail, twice during the growing period.

For the RNA-sequencing there were three soil Zn treatments: no Zn addition, 5 mg Zn kg^-1^, and 20 mg Zn kg^-1^. Plants were destructively harvested after 33 days, and roots washed clean with RO water were snap frozen in liquid nitrogen before storage at −80 °C.

For the other plant growth experiments, plants were destructively harvested after 35 days. Shoots were separated at the soil level and roots were washed clean before a subsample of fresh root biomass was moved into 70 % ethanol for determination of AM colonisation. The shoots and remaining root material were dried at 60 °C for at least 48 hours before dry weights were determined. Following that, the entire shoot material was homogenised and digested in 4:1 nitric acid:hydrogen peroxide at 125 °C for three hours before being diluted with RO water and analysed for elemental concentrations of P, Mg, K, Zn, Mn, and Fe by ICP-OES. The fresh root subsamples were rinsed well and moved into a 10% sodium hydroxide solution at room temperature for seven days to clear the root cells. Cleared roots were rinsed well then stained in a 5 % ink in vinegar solution (34) at 60 °C for 10 minutes before being stored in 50 % glycerol. Colonisation by *R. irregularis* was determined on the stained roots following (35) whereby arbuscular, vesicular and hyphal root length colonised were each independently estimated on 100 roots intersects per sample.

## RNA sequencing

### RNA isolation and sequencing

For all experiments, a subsample (∼100 mg) flash frozen root material was homogenised in 2 mL microcentrifuge tubes with two 2.8 mm ceramic beads per tube, in a bead beater for 2 × 30 seconds (Genogrinder). Total RNA was subsequently isolated using a Plant Total RNA kit (Sigma) including on-column DNase treatment following the manufacturer’s instructions. The quality and yield of the resulting RNA was analysed using a BioAnalyzer instrument (RNA-sequencing) or Nandrop (qRT-PCR). Three biological replicates of each treatment were used in the library preparation for RNA sequencing. RNA-seq libraries were prepared using the TruSeq Stranded mRNA Library Prep Kit according to the manufacturer’s instructions (Illumina) and sequenced on a NextSeq 550 system (Illumina) as 75bp single-end reads with an average quality score (Q30) of above 92%. RNA-seq data was deposited at the NCBI Sequence Read Archive (NCBI SRA) under project ID PRJNA660297.

### Bioinformatics and analysis of differentially expressed genes

Quality control of RNA-seq data was performed using the FastQC software (https://www.bioinformatics.babraham.ac.uk/projects/fastqc/). Transcript abundances as transcripts per million (TPM) and estimated counts were quantified on a gene level by pseudo-aligning reads against a k-mer index build from the representative transcript models downloaded for the *Medicago truncatula* Mt4.0 annotation using a k-mer length of 31 (36) using the kallisto program with 100 bootstraps (37). Only genes with at least five counts were included in the further analysis. The program sleuth with a Wald test was used to test for differential gene expression (38). Differentially expressed genes (DEGs) were calculated as the log fold change (FC) of the mean *R. irregularis*-inoculated plants to the mock-inoculated plants, for each soil Zn treatment, respectively. Genes were considered as differentially expressed with a |log_2_ (fold change)| > 2 and false discovery rate (FDR) < 0.05.

For further analyses, hierarchical clustering and generation of heat maps the Partek Genomics software suite version 6.16 (Partek Incorporated, http://www.partek.com/) was used. Venn diagrams were constructed (http://bioinformatics.psb.ugent.be/webtools/Venn/) to visualise the separation of DEGs into the three Zn treatments. GO term enrichment analysis of the DEGs was completed using the agriGO tool (http://bioinfo.cau.edu.cn/agriGO/analysis.php).

## Characterisation of *MtZIP14*

### Expression of MtZIP14 in roots colonised by AM fungi

To confirm the expression pattern of *MtZIP14* in AM-inoculated compared with mock-inoculated plants, qRT-PCR was performed on material from an independently conducted experiment (see 39). Briefly, *Medicago truncatula* A17 was inoculated with *R. irregularis* or mock-inoculated and grown in a soil with one of four different soil Zn concentrations. Total root RNA was isolated as described above and expression of *MtZIP14* was measured by qRT-PCR and normalised to the geometric mean of three housekeeping genes following (40) (Table S5).

### Localisation of MtZIP14 in planta

A DNA fragment of the full-length MtZIP14 (Medtr6g007687.1) genomic region and 1.9 kb upstream of the ATG, but without the stop codon or 3’ UTR, was cloned into the pENTR-D-TOPO vector as above, confirmed by Sanger sequencing, and recombined into the Gateway-compatible C-terminal GFP-tagged plant expression vector pMDC107 by LR reaction. Successful recombination into the expression vector was confirmed by enzyme digestion and Sanger sequencing. The resulting pMtZIP14:MtZIP14-GFP construct and the empty pMDC107 vector were respectively transformed in *Agrobacterium rhizogenes* ARqua1, confirmed by colony PCR, and used for hairy root transformation of *M. truncatula* cv. A17 plants.

Hairy roots of *M. truncatula* A17 plants were transformed with the constructs (MtZIP14-GFP and empty vector, respectively) following Floss, Schmitz, Starker, Gantt and Harrison (29) and grown on F-media containing 25 µg mL^-1^ kanamycin for three weeks. Seedlings with roots developing from the inoculation site were then moved into sterilised zeolite substrate for 10 days and finally to the soil/sand substrate inoculated with *R. irregularis* (described above).

At 21 to 26 days post-inoculation with *R. irregularis* the plants were gently removed from the substrate and roots washed free of any sand/soil. Roots were examined under a Nikon SMZ25 stereomicroscope with a 2× objective for evidence of AM colonisation (external hyphal penetration, swelling) sectioned transversely into 1-2 mm pieces, then longitudinally, before mounting onto slides for further viewing on the confocal microscope. Images were captured with a Nikon A1R Confocal Laser-Scanning Microscope, using the 60× Plan Apo VC WI objective; excitation 488 nm, emission collection 500-550 nm; an image using the transmission detector was simultaneously captured.

Three independent hairy root transformation events were conducted over a five-month period, with at least six plants assessed from each event. The images presented are representative of the GFP and empty vector constructs, respectively.

### Complementation of a Zn-deficient yeast strain with MtZIP14

The complete mRNA sequence of MtZIP14 cv. A17 from ATG to stop codon was amplified using Phusion High-fidelity DNA polymerase, with the Gateway-specific sequence (CACC) added to the 5’ end of the forward primer. The resulting product was cloned into the pENTR-D-TOPO Gateway-compatible entry vector, transformed by heat shock into *E. coli* DH5α competent cells, and sequenced to confirm before LR reaction to recombine into the yeast expression vector pDEST52. Successful recombination was confirmed by enzyme digestion. Then, the pDEST52:MtZIP14 construct and empty pDEST52 vector were respectively transformed into the yeast strain ZHY3 (*zrt1zrt2* mutated) and the wild-type yeast strain DY1457 (both kindly provided by Prof. D. Eide) using the lithium-acetate transformation method (41). Transformants were selected on YNB minus uracil plus 2 % glucose plates.

For the yeast growth studies, the ZHY3 yeast strain expressing empty vector and the MtZIP14 constructs were grown overnight in liquid yeast nitrogen base (YNB) -uracil with 2 % galactose. Cells were pelleted, washed three times in sterile water and resuspended in YNB -uracil with 2 % galactose media supplemented with 1 mM EDTA and one of 0.1, 0.2, 0.5, 1.0 or 1.5 mM Zn as ZnSO_4_ to an OD of ∼0.23. The EDTA was added for the purpose of chelating the existing Zn in the media (400 µg Zn L^-1^ according to manufacturer) and allowed for the creation of media completely devoid of Zn (following 12). The availability of free Zn^2+^ in the EDTA-YNB media was predicted using the Visual MINTEQ software (https://vminteq.lwr.kth.se/). Without any addition of ZnSO_4_, the EDTA-YNB media was predicted to have approximately 0.001 nM free Zn^2+^, and the addition of 0.1 mM ZnSO_4_ yielded just 0.048 nM free Zn^2+^. At the highest ZnSO_4_ addition of 1.5 mM, the EDTA-YNB media had a predicted free Zn^2+^ availability of 237.86 µM.

An aliquot of 150 µl of cells was placed into a 96 well microplate for each treatment (3 replicates of each) and the plate sealed with a sterile film. The microplate was placed into a microplate reader (BMG Omega) and growth of the yeast strains was quantified over 66 h. Solid agar plates were prepared from YNB -uracil with 2% galactose and 1 mM EDTA with the addition of Zn at 0, 0.2, 0.5 or 1 mM ZnSO_4_. The wild-type and ZHY3 yeast empty vector constructs and the ZHY3-MtZIP14 construct were cultured overnight in 5 mL of YNB -uracil with 2% galactose. The resulting cultures were rinsed well with sterile water three times before being resuspended in 3 mL sterile water and diluted to an OD_600_ of 0.5, 0.1, 0.01 and 0.001. For each yeast construct, 5 µL of each dilution was spotted onto the prepared EDTA-YNB agar plates, and onto control YNB -uracil with 2% galactose or 2% glucose (no EDTA) plates and placed inverted in a 28 °C incubator for 2-4 days. The experimental plates were replicated three times. The plates with wild-type yeast harbouring the empty vector construct were photographed after two days and the ZHY3 yeast harbouring the empty vector or MtZIP14 constructs after four days, due to faster growth of the WT strain.

### Phenotyping of loss-of-function MtZIP14 mutants

To find the *MtZIP14* gene sequence in the *M. truncatula* R108 ecotype, the *MtZIP14* A17 ecotype mRNA sequence was compared using a BLAST online tool (http://www.medicagohapmap.org/tools/r108_blastform).

Line numbers NF8057 and NF4665 from the Noble Foundation’s *M. truncatula* Tnt1 insertion mutant collection were predicted to have an insertion in the *MtZIP14* gene as per BLAST analysis of R108 sequence in the Tnt1 insertion collection database (https://medicago-mutant.noble.org/mutant/database.php). The NF8057 line (referred to in text and figures as “knock-out”, KO) has a Tnt1 insertion in the first exon of the MtZIP14 gene sequence, approximately 250 nucleotides downstream of the ATG. The NF4665 line (referred to in text and figures as “knock-down”, KD) has a Tnt1 insertion in the first exon approximately 496 nucleotides downstream of the ATG. Genotyping of the R1 plants supplied by the Noble Foundation using gene-specific and Tnt1-specific primers identified a number of plants homozygous for the Tnt1 insertion, which were subsequently genetically backcrossed using the keel petal incision method to the R108 wild-type (WT) background (following 42). The resulting heterozygous progeny were grown and allowed to self-pollinate, then homozygous and out-segregated WT progeny were isolated for use in subsequent experiments.

To investigate the *mtzip14* phenotype, *M. truncatula* R108 *mtzip14*, the respective out-segregated WT for each NF line, and R108 wild-type plants were inoculated with *R. irregularis* or with mock inoculum and grown in a Zn-deficient soil, as described above. Each treatment was biologically replicated six times. Plants were harvested 35 days after transplantation; measurements of dry shoot and root biomass, AM colonisation (arbuscular, vesicular, hyphal), and shoot Zn concentration were taken. Flash frozen root samples were taken from one experiment for the isolation of RNA and gene expression analysis by qRT-PCR (oligo sequences in Table S5). The phenotyping experiment was conducted in an identical manner a total of three times over four months.

### Statistical analysis and data presentation

A linear mixed effects model was employed to analyse the plant biomass, Zn nutrition and AM colonisation data using the “lme” function within the “nlme” package in R version 4.0.2 (R Core Team 2019). *Mycorrhiza* and *Genotype* (and their interaction) were included as fixed effects and a random term for *Experiment* was included in order to block the data by the experiment it originated from. This allowed for data from all three growth experiments to be included in the model while accounting for effects of the individual *Experiment*. The NF4665 (knock-down) and NF8057 (knock-out) Tnt1 lines were statistically analysed separately. For the NF8057 gene expression data collected from one experiment, a two-way analysis of variance (ANOVA) was employed with *Mycorrhiza* and *Genotype* as the factors. Where the interaction or main effects was significant (*P*<0.05), the “lsmeans” package and function were used to conduct Tukey’s HSD *post hoc* pairwise comparisons between the treatments and identify any significant differences. These are presented as letters on the relevant figures.

The Tnt1 plant physiological and gene expression data are presented as box-and-whisker plots (one representative experiment presented in main figures, boxplots from all three experiment available in Figure S8a-f), and were generated using the “ggboxplot” function within the “ggpubr” package with “jitter” added to visualise the individual data points and outliers. A principal components analysis (PCA) was undertaken using the “PCA” function in the “FactoMineR” package, including all of the available plant biomass, AM colonisation and nutrient content data to visualise the effect of *Mycorrhiza* and *Genotype* on the data. Following that, the AMF and Mock data were split, and PCA conducted on each dataset separately to visualise the effect of *Genotype*. The PCA biplots were drawn using the “factoextra” package and the scores coloured by levels of *Mycorrhiza* (AMF or Mock) or *Genotype* (*mtzip1*4 or WT); the group mean was also computed for each level, and a 95 % confidence ellipse drawn around the mean to determine significant differences between groups.

## Acknowledgements

SJWW acknowledges the University of Adelaide Ramsay Fellowship and the Australasian Mycological Society research grant for support. We thank Gwen Mayo and Adelaide Microscopy for support with imaging. All authors acknowledge the Australian Research Council Centre of Excellence in Plant Energy Biology (CE1400008) for support. We wish to thank Dr Armando Bravo, Prof. Maria Harrison, Prof. Timothy Cavagnaro, Dr Apriadi Situmorang, and Dr Yoshihiro Kobae for valuable discussions, Prof. David Eide for access to the yeast strains and Dr Jianqi Wen for access to the *M. truncatula* Tnt1 insertion mutant collection. We would like to thank Asha Haslem for technical assistance and the La Trobe University Genomics Platform for access to next-generation sequencing equipment.

The *Medicago truncatula* plants utilized in this research project, which are jointly owned by the Centre National De La Recherche Scientifique, were obtained from Noble Research Institute, LLC (successor-by-conversion to The Samuel Roberts Noble Foundation, Inc., effective May 1, 2017) and were created through research funded, in part, by a grant from the National Science Foundation, NSF-0703285.

