## Supplementary Tables 5-8; Supplementary Figures 1-8 for "Identification of a unique ZIP transporter involved in zinc uptake via the arbuscular mycorrhizal fungal pathway"

**Table S5.** Forward (F) and reverse (R) oligo sequences used for the amplification of genes in the *Medicago truncatula* A17 and R108 ecotypes by quantitative RT-PCR.

| Name | Phytozome ID | Sequence |
|---|---|---|
| MtEF1a-F (housekeeping) | Medtr6g021800 | TGACAGGCGATCTGGTAAGG |
| MtEF1a-R | | TCAGCGAAGGTCTCAACCAC |
| MtASPP – F (housekeeping) | Medtr4g095270 | GGATCGGTCTTGGACAGTGG |
| MtASPP – R | | TGGACCGCTGATTTGACTGA |
| MtTRF – F (housekeeping) | Medtr1g087790 | GGATAAGGTGGATGGTGATCG |
| MtTRF – R | | TCTGCCTCTCGTCGTTTTTGT |
| *Medtr6g051680* – F (HMA) | Medtr6g051680 | GGCTTGATGCTGTTTCAATCG |
| *Medtr6g051680* - R | | ACTTCGACACAACACTAACAGGA |
| *Medtr0041s0140* – F (HMA) | Medtr0041s0140 | AGGTAGAGGTTAATAAAATGGAGCA |
| *Medtr0041s0140* - R | | GGTTCATATCCGTCATACCAATACA |
| *Medtr8g035880* – F (Zn-binding) | Medtr3g089970 | CCAAGTTCATGCTGCTGCTTT |
| *Medtr8g035880* - R | | CGGGAAACTCAGACGGAAAA |
| MtZIP14 - F | Medtr6g007687 | GCATCTGCAGGGGTTCTCAT |
| MtZIP14 - R | | AAGTGCTAAACTTGCCCCGA |
| MtZIP1 - F | Medtr2g064310 | GGGATAGCAATTGGGATGGGA |
| MtZIP1 - R | | GCAAGTAGGTCAACTAGTGCCA |
| MtZIP2 - F | Medtr2g097580 | AATGGGCATTGCTTGTGGTG |
| MtZIP2 - R | | TGTCGAAACGGCTCTTCCTC |
| MtZIP3 - F | Medtr3g081580 | TGGCATAGGGATCAGTAGTGG |
| MtZIP3 - R | | CTACCGCTCTTTTGCATCCTTG |
| MtZIP4/10 - F | Medtr3g082050/ | GGACAGATCAAAAGCTCTACGC |
| MtZIP4/10 - R | Medtr3g081690 | TCACTTTCCCAAGAAGTGGGAG |
| MtZIP5 - F | Medtr1g016120 | ACGTGTGGTCAGAGTTTCCA |
| MtZIP5 - R | | AACAACTCGTCCGTGCTCAG |
| MtZIP6 - F | Medtr4g083570 | CTTGGCGACACGTTCAATCC |
| MtZIP6 - R | | CATGAACCCGGTCCCAAGAA |
| MtZIP7 - F | Medtr3g058630 | CGCCCTTTGTGCTCATTCTG |
| MtZIP7 - R | | GCTTTCCATGCGTCTGCTTT |
| MtZIP9 - F | Medtr3g081640 | TGGCAAAGTAATACCGGCGT |
| MtZIP9 - R | | CACCGGCAGCAAAGGTTTTA |
| MtZIP11 - F | Medtr3g104400 | TTGGGCTGTCTTTGGGAGTT |
| MtZIP11 - R | | CCCTCCAAGAGCAAACCCTT |
| MtZIP15 - F | Medtr7g074060 | ACGCTCTTCTCATGGGCATT |
| MtZIP15 - R | | AGCAGCTCCCAATGCAAGAA |
| MtPT4 - F | Medtr1g028600 | GACACGAGGCGCTTTCATAGCAGC |
| MtPT4 - R | | GTCATCGCAGCTGGAACAGCACCG |

**Table S6.** Statistical outcomes (*P*-values) from the two-way analysis of variance conducted on NF8057 gene expression data. The highest order significant term/s are highlighted in bold.

| | *Myc* | *Genotype* | *Myc*Genotype* |
|---|---|---|---|
| *MtZIP1* expression | **0.0003** | 0.4281 | 0.462 |
| *MtZIP2* expression | **<0.0001** | 0.3528 | 0.3926 |
| *MtZIP3* expression | **<0.0001** | 0.9058 | 0.9731 |
| *MtZIP4/10* expression | **<0.0001** | 0.5008 | 0.4811 |
| *MtZIP5* expression | **<0.0001** | 0.3096 | 0.7705 |
| *MtZIP6* expression | **0.0001** | 0.7778 | 0.1922 |
| *MtZIP7* expression | **<0.0001** | 0.0797 | 0.1870 |
| *MtZIP8* expression | <0.0001 | 0.0286 | **0.0321** |
| *MtZIP9* expression | 0.071 | 0.6227 | 0.7037 |
| *MtZIP11* expression | **<0.0001** | 0.3181 | 0.8250 |
| *MtZIP12* expression | 0.0644 | 0.4715 | 0.2129 |
| *MtZIP15* expression | **0.0275** | 0.1722 | 0.3045 |

**Table S7.** Statistical outcomes (*P*-values) from the linear mixed effects model for NF4665. The highest order significant term/s are highlighted in bold.

| | *Myc* | *Genotype* | *Myc*Genotype* |
|---|---|---|---|
| Shoot dry weight | 0.0001 | 0.0791 | **0.0425** |
| Shoot Zn content | <0.0001 | 0.1606 | **0.0197** |
| Shoot Zn concentration | **<0.0001** | 0.637 | 0.827 |
| Arbuscular colonisation | <0.0001 | 0.0129 | **0.0103** |
| Vesicular colonisation | <0.0001 | <0.0001 | **<0.0001** |
| Hyphal colonisation | **<0.0001** | 0.1489 | 0.1363 |
| *MtZIP14* expression | <0.0001 | <0.0001 | **<0.0001** |

**Table S8.** Statistical outcomes (*P*-values) from the linear mixed effects model for NF8057. The highest order significant term/s are highlighted in bold.

| | *Myc* | *Genotype* | *Myc*Genotype* |
|---|---|---|---|
| Shoot dry weight | 0.0015 | 0.0003 | **0.0037** |
| Shoot Zn content | <0.0001 | 0.024 | **0.0334** |
| Shoot Zn concentration | **<0.0001** | 0.8244 | 0.9807 |
| Arbuscular colonisation | <0.0001 | 0.0016 | **0.0016** |
| Vesicular colonisation | <0.0001 | <0.0001 | **<0.0001** |
| Hyphal colonisation | **<0.0001** | 0.0889 | **0.0843** |
| *MtZIP14* expression | <0.0001 | <0.0001 | **<0.0001** |

a

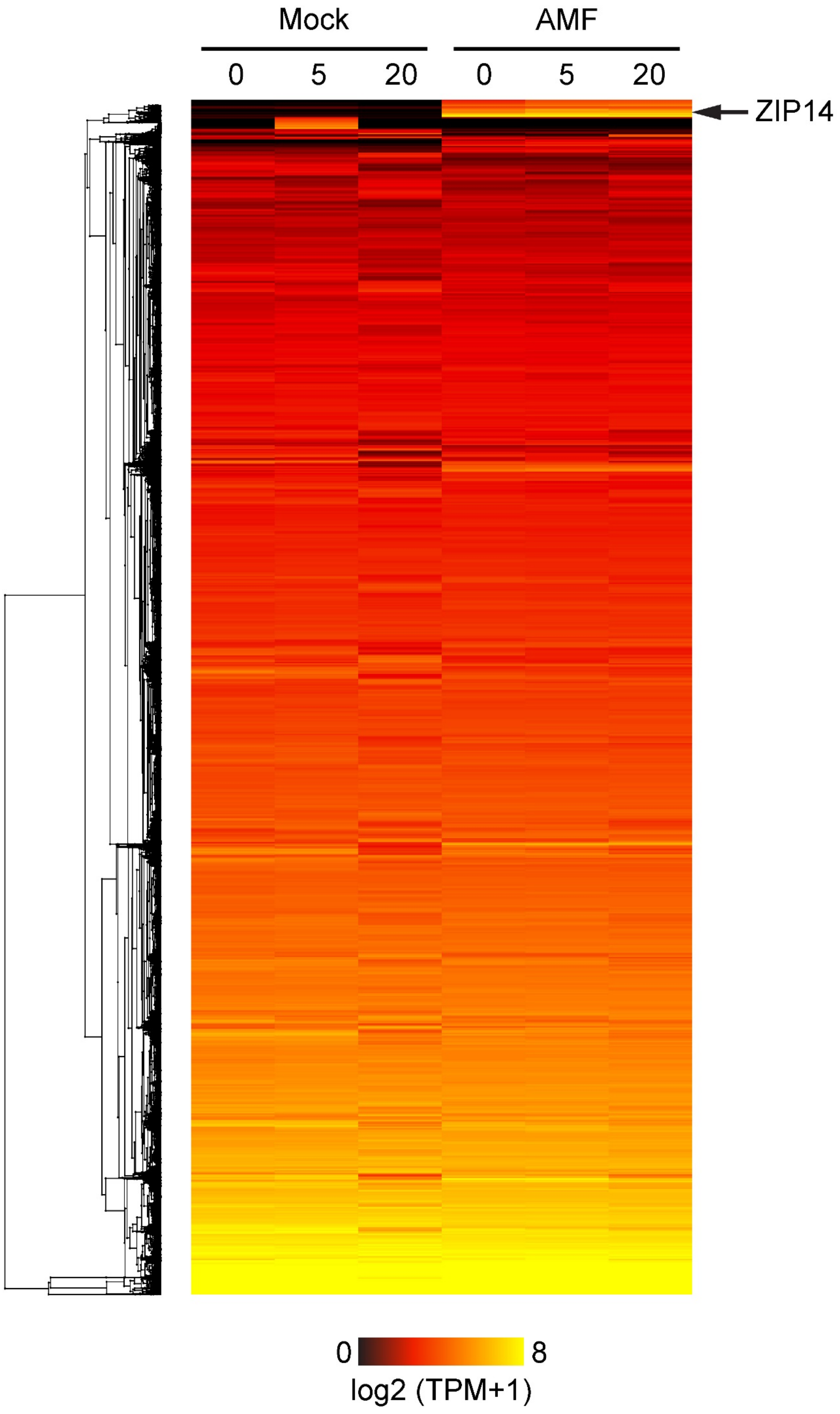

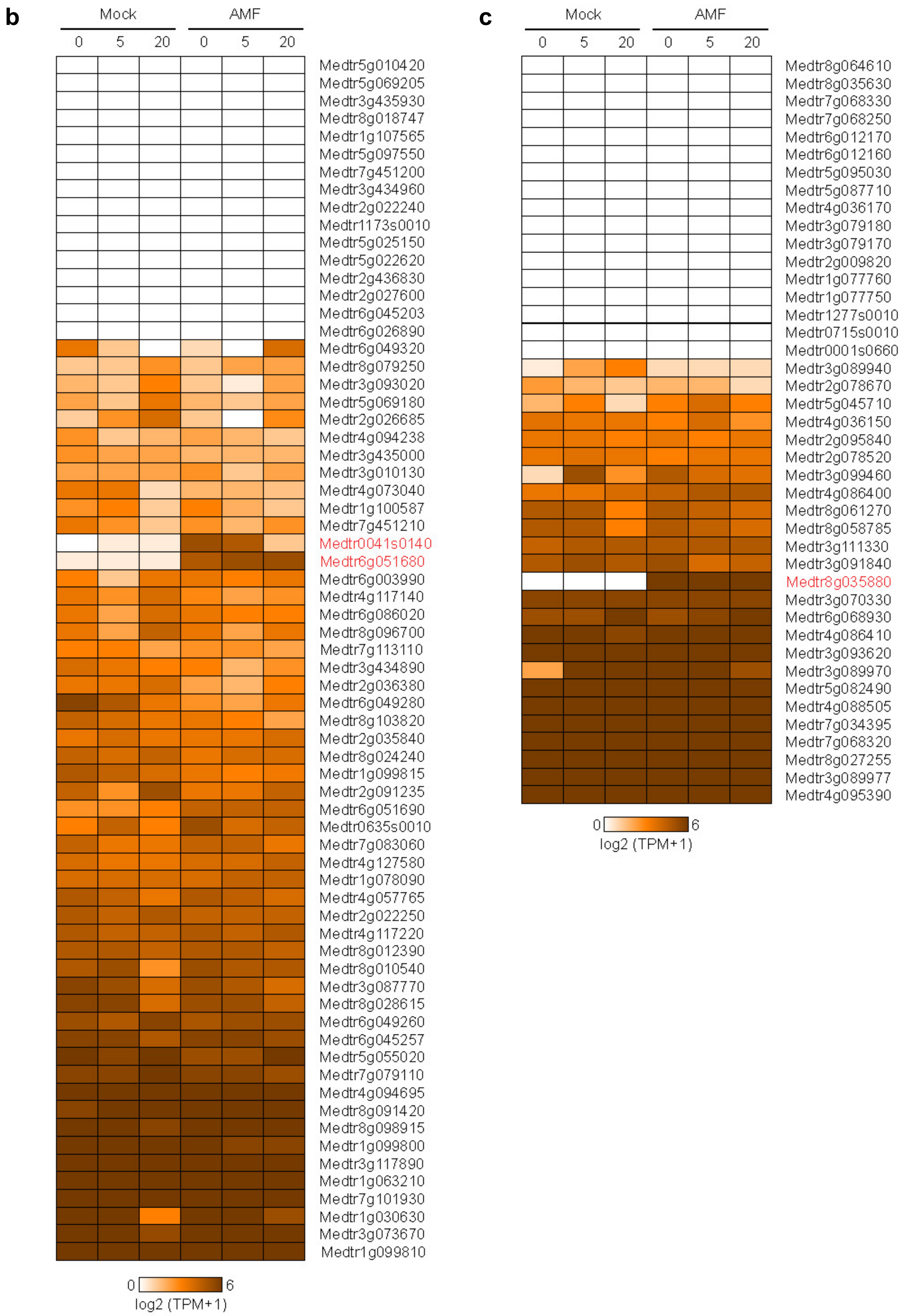


**Figure S1.** Heat maps from the emtire *Medicago truncatula* root transcriptome (see Table S1) (a) and filtered lists for genes with the annotation including the term heavy metal transport (b) or zinc-binding (c), split into three soil Zn addition treatments: Zn0 no addition; Zn5 5 mg $kg^{-1}$ addition; Zn20 20 mg $kg^{-1}$ addition. Genes highly up-regulated in AM colonised plants across all three Zn addition treatments are highlighted in red.

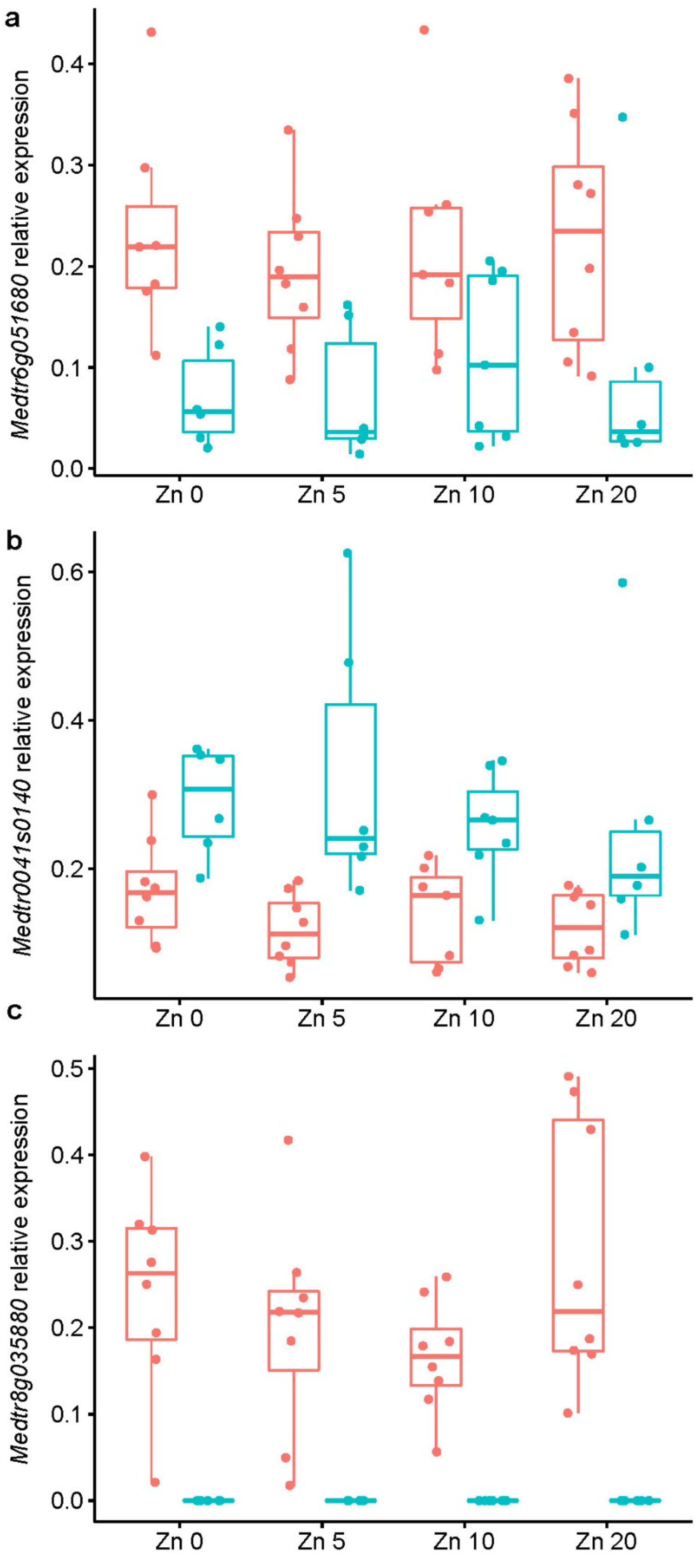


**Figure S2.** Expression of two *Medicago truncatula* heavy metal associated (HMA) domain protein genes (a-b) and a gene encoding for Zn-binding dehydrogenase oxidoreductase (c) in the roots that were *Rhizophagus irregularis*-inoculated (pink) and mock-inoculated (blue) plants grown at four soil Zn additions. Gene expression is calculated as the gene-of-interest relative to the geometric mean of three housekeeping genes (Table S5).

```
CLUSTAL O(1.2.4) multiple sequence alignment


MtZIP2       MASFKT---LKSTFLILCLLAS--FFLNPIKAHGGHDDSDHDSDNINIRSRSLVLVKIWC      55
MtZIP5       ------------------MSTLFDEFMTNSSCESGESDLCRDE------S-AALILKFVA      35
MtZIP14      MIKFVTSKFMTISIII-LLQQNLVFS--KCSCENQVEDSYHKV------S-EALKYKLIA      50
MtZIP1       -MARLNNPSILVTIIL-FLLVTLPFASCESECSSKYEGVCHNK------N-EALKLKLIA      51
MtZIP6       -MAY-SVTIHKAIFIV-FILITFLTSQALADCESESTNSCNNK------E-KAQPLKLIA      50
                               :           .. .   .  ..       .      *: .

MtZIP2       LIILFVFTFIGGVSPY------YFRWNEVFLLLGTQFAGGVFLGTSMMHFLSDSNETFED      109
MtZIP5       MASILVAGFSGIAVPLLGNRRGLLRSDGEILPAAKAFAAGVILATGFVHMLQDAWKALNH      95
MtZIP14      MATVFVSSLIGVCIPIFAKKCSYLNPENDFYFLVKAFAAGVILATGFIHILPDAFEALTS      110
MtZIP1       IFSILVTSMIGICIPIFTTSIPALKPDGDLFVIIKAFASGVILATGYMHVMPDSFQDLNS      111
MtZIP6       IFSILATSVIGVCLPLATRSIPALSPEGDLFIIVKCFAAGIILGTGFMHVLPDSYEMLWS      110
             :  ::.  . *   *        :  :  :    . **.*::*.*. :*.: *: : :

MtZIP2       LTK-------KTYPFAFMLACSGYLLTMFGDCVVV-YVTSNNQREAKVE--------ELE      153
MtZIP5       SCLKSYSHVWSEFPFTGFFAMMSALLTLLVDFVATQYYESQHQKTHDRHGRVVGNGEGLE      155
MtZIP14      PCISE--KPWKLFPFSGFVTMVAAIGTLIMEALIMGYHKRSEM-KKAQP---------LD      158
MtZIP1       PCLPE--RPWKKFPFTTFIAMVSAVFTLMVDSFSISFFKKKLSASSSSN---------LE      160
MtZIP6       DCLDE--KPWHEFPFSGLVAMFSAVVTMMVDSIATSYYSKKGKSGVVIP-----------      157
                         :**: :.:  . : *:: : .   :   .

MtZIP2       -------------------------------GGRTPQEE-EGTT--------ELAMDESN      173
MtZIP5       EELLGSGIVEVQGETFGGGMHIVGMHAHASQHGHSHQNHGDGHG-HGHSHSFG---EHDG      211
MtZIP14      ----------------------------------ENDETHHSDNGSSHVHNFSIASDRLD      184
MtZIP1       ----------------------------AGSETKEPEQIGHGHGHGLVVAN-GHE-KNVN      190
MtZIP6       -----------------------------ESHGGDDQEIGHSHGGHHHIHN-GFKTEESD      187
                                                 ::  ..              .. .

MtZIP2       VAFMKTTNVGDTILLILALCFHSVFEGIAVGISGTKEEAWRNLWTISLHKIFAAIAMGIA      233
MtZIP5       VDSS-VRHVVVSQVLELGIVSHSLIIGLSLGVSQSPCTMRPLIAALSFHQFFEGFALGGC      270
MtZIP14      STNR-LRYTIVSQILELGIVLHSVILGISLGVSRSPKTIKPLVAVLTFHQCFEGIGLGGC      243
MtZIP1       AEQL-MRYRVVAQVLELGIVVHSVVIGLSLGASENHCTIRPLIAALCFHQLFEGMGLGGC      249
MtZIP6       EPQL-LRYRVVVMVLELGIVVHSVVIGLGMGASNNTCSIKGLIAAMCFHQMFEGMGLGGC      246
                          :* *.:  **:. *:.:* * .       : .: :*: * .:.:* .

MtZIP2       LLRMLPKRPLITTAGYSFAFAISSPIGVGIGIAIDATTEGKTA--DWMYAISMGIACGVF      291
MtZIP5       ISEARFKTS--SATIMACFFALTTPLGVAIGTLVASNFNPYSPGALITEGILDSLSAGIL      328
MtZIP14      ISQAQFKYY--KVTIMILFFCLIFPIGIGIGMGISNIYNESSPKSLIVEGFLLSASAGVL      301
MtZIP1       ILQADYGTK--MKSTMIFFFSATTPFGIALGIGLSKVYSNTSPTALIVEGVLNAMSAGLL      307
MtZIP6       ILQAKYKFL--KNAMLVFFFSITTPLGIAIGLAMSTSYKENSPVALITVGLLNASSAGLL      304
             : .          :     *.   *:*:.:*  :    .  :       ..  . :.*::

MtZIP2       VYVAINHLISKGFKPQR-KSRFDTPWFRFLAVLFGVAVIAVVMIWD*-------------      336
MtZIP5       VYMALVDLIAADFLSKKMRCSLRLQIVSFCLLFLGAGSMSSLALWA*-------------      374
MtZIP14      INMALVDLVATDFMNSKMLTNFRLQLGASLALFVGMICMSILALGEDS*-----------      349
MtZIP1       NYMALVDLLANDFMGAKLQSRMKLQIWSYVAVLLGAGGMSDAFHHRHVASEDDEDEEQVF      367
MtZIP6       IYMALVDLLAADFMSKRMQSSIKLQLKSYVAVFLGAGGMSLMAKWA*-------------      350
               :*: .*:: .*   :    :         ::.*   ::

MtZIP2       ------------------------------     336
MtZIP5       ------------------------------     374
MtZIP14      ------------------------------     349
MtZIP1       VPSPLLSLKEQIEKDKGRGEAHDELRIEI*     396
MtZIP6       ------------------------------     350
```

**Figure S3.** Amino acid alignment of MtZIP14 with four other *Medicago truncatula* ZIP transporters (b) that have been confirmed to transport Zn in the same *zrt1zrt2* yeast system. A potential metal-binding region rich in histidine can be observed between transmembrane III and IV (beginning AA 164 in MtZIP14), as well as the proposed conserved ZIP signature domain (beginning AA 194 for MtZIP14).

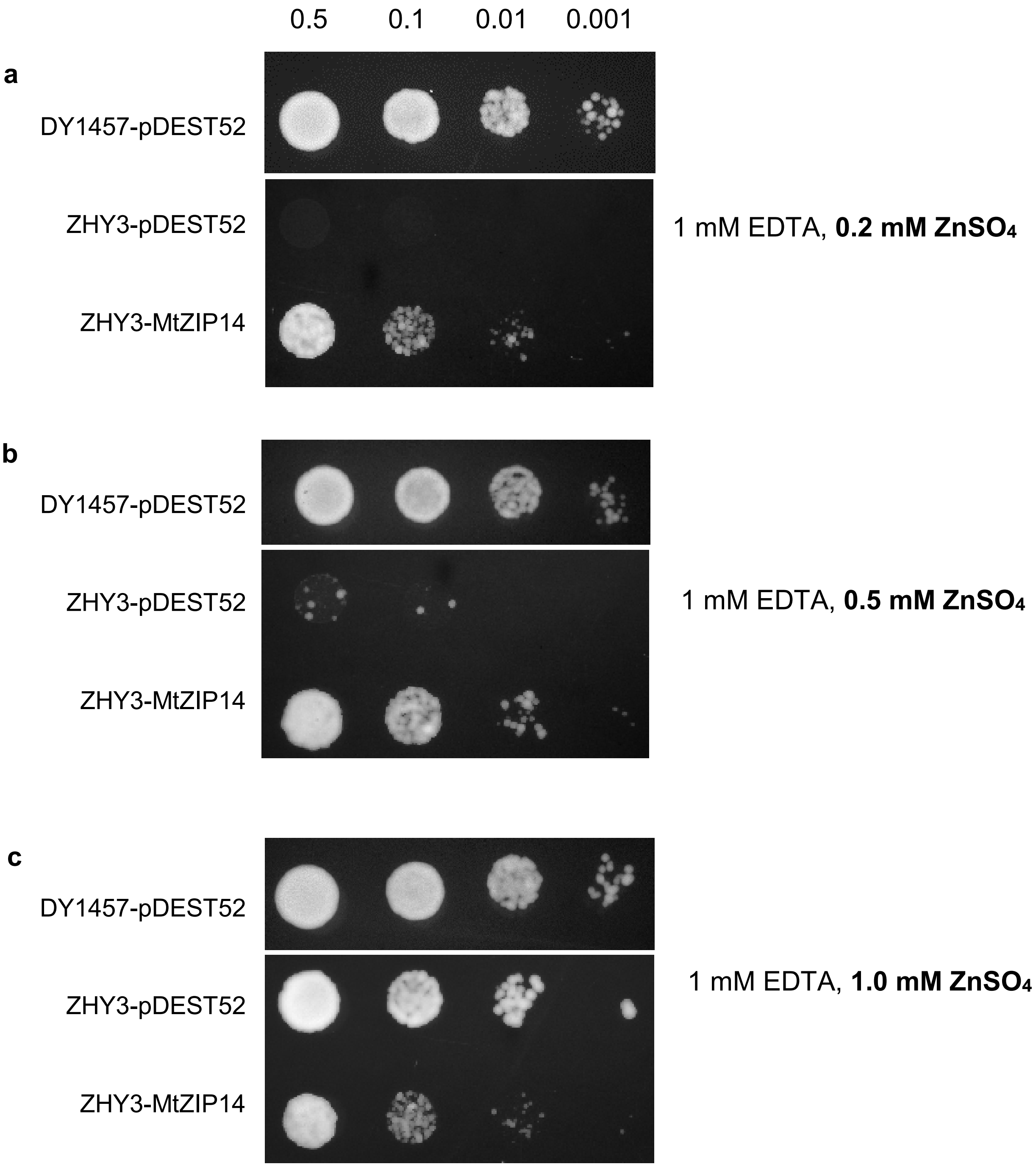


**Figure S4.** Yeast Zn transporter (*zrt1zrt2*) mutant (ZHY3) and wild-type (DY1457) strains expressing MtZIP14, or the empty vector pDEST52, grown on solid YNB -uracil media with 2% galactose and 1mM EDTA. With the addition of EDTA, there was no growth of any yeast strains without Zn supplementation. The Zn supplementation treatments were 0.2 (a), 0.5 (b) and 1.0 mM (c) $ZnSO_4$ and dilutions of the yeast were spotted at 0.5, 0.1, 0.01 and 0.001 $OD_{600}$. Due to the faster growth of the wild-type strain, DY1457 was photographed after 2 days incubating at 28 °C and ZHY3 was photographed after 4 days.

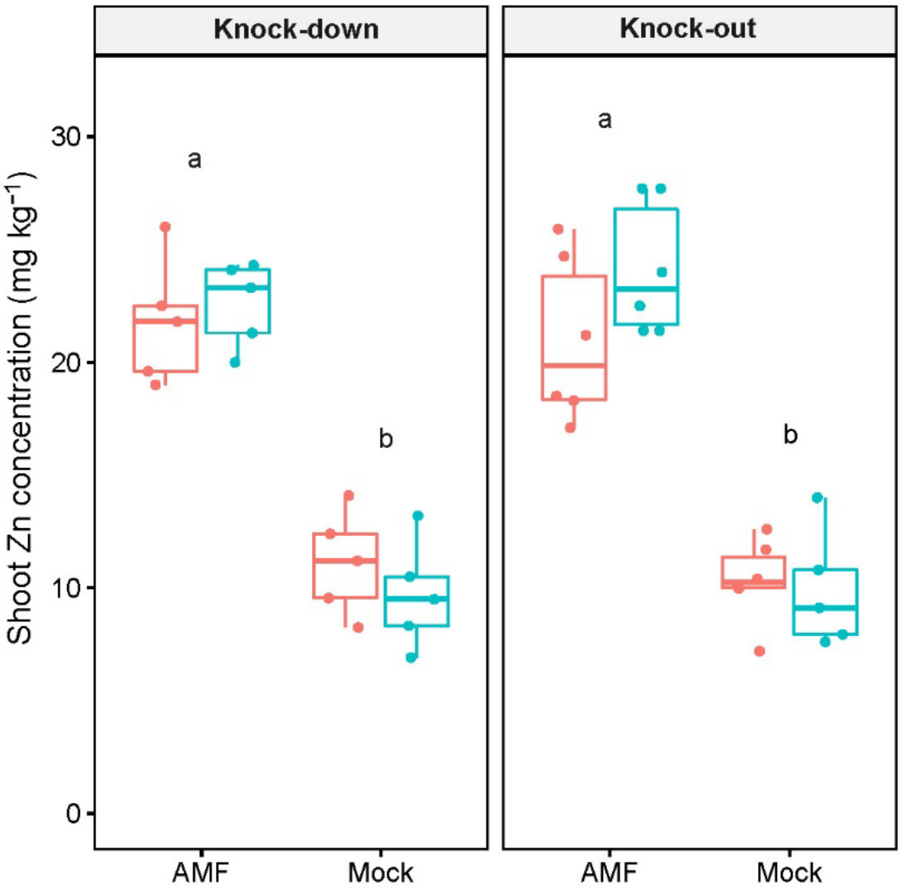


**Figure S5.** Shoot Zn concentration in the two *mtzip14* (pink) and segregating WT (blue) lines grown with or without inoculation by the AM fungus *Rhizophagus irregularis*. Means with different letters are considered significantly different ($P$<0.05) as per Tukey's HSD *post hoc* test. Where one letter appears above two boxes, it represents a significant main effect of *Mycorrhiza* where the two genotypes are pooled.

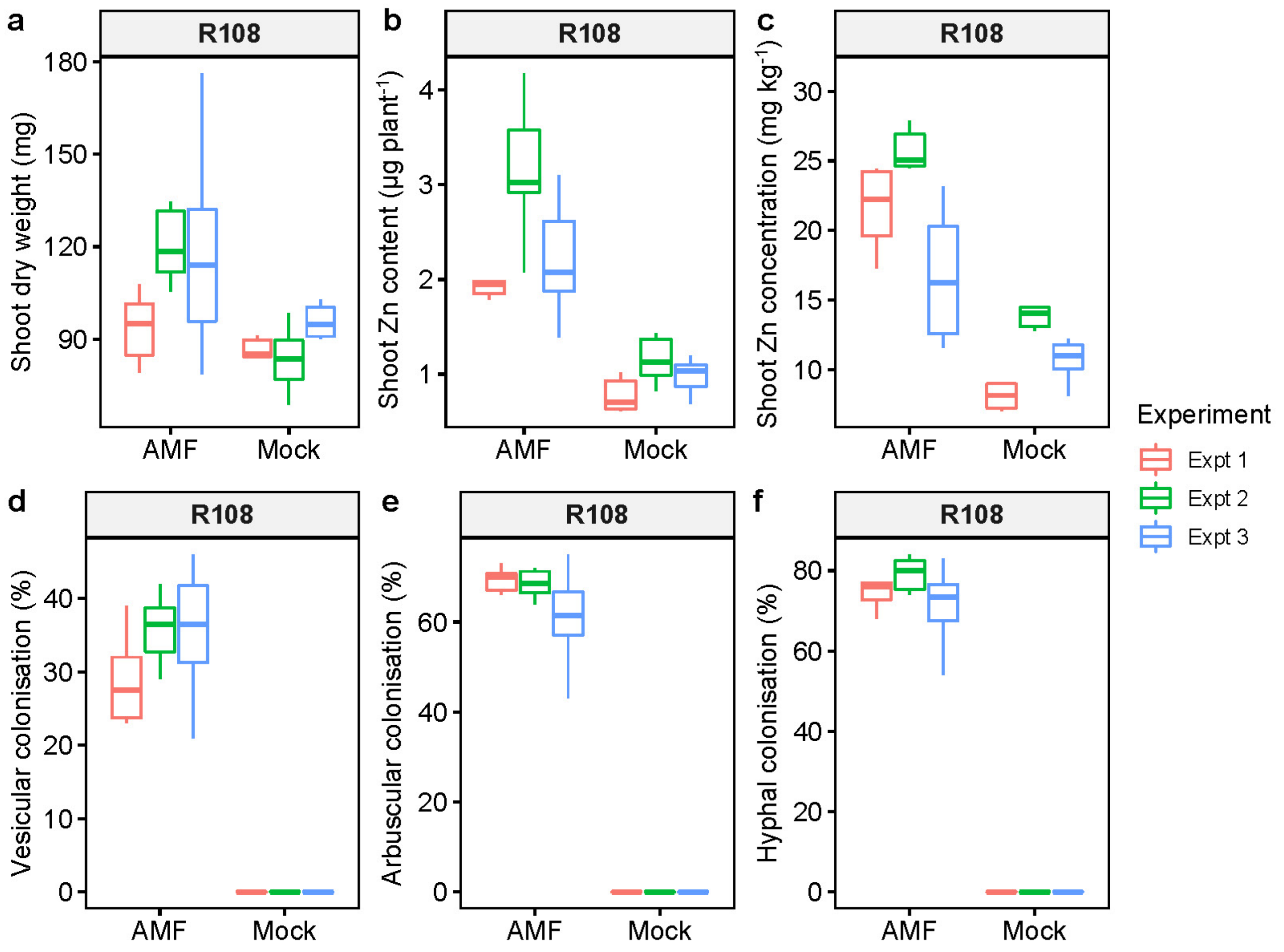


**Figure S6.** Plant biomass (a), zinc nutrition (b-c) and arbuscular mycorrhizal colonisation (d-f) data from the *Medicago truncatula* R108 wild-type plants grown with the Tnt1 lines in three independent experiments, for comparison with the segregating wild-type data.

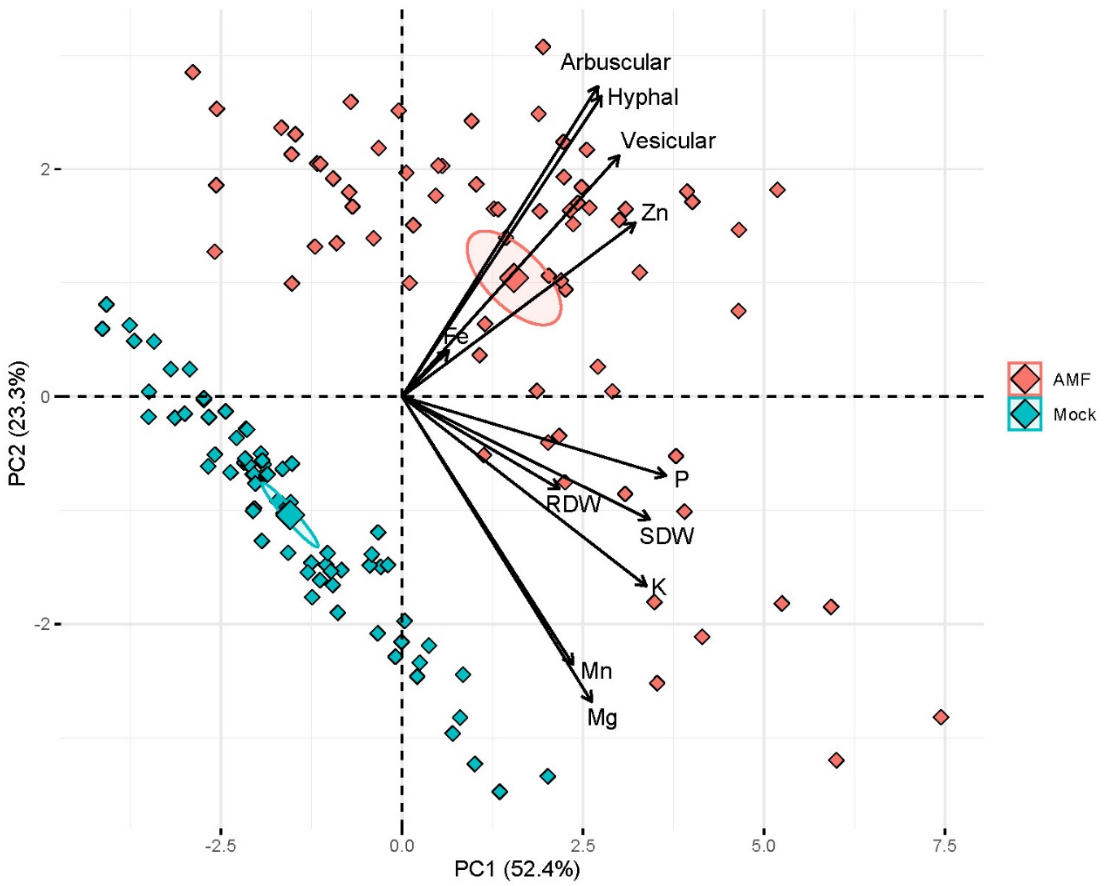


**Figure S7.** Principal components analysis (PCA) biplot displaying scores in the first two principal components (PC1: x-axis, PC2: y-axis) following PCA of biomass (shoot and root dry weights), shoot nutrient contents (µg plant$^{-1}$; Zn, P, Mg, K, Mn, Fe) and arbuscular mycorrhizal colonisation (arbuscular, vesicular, hyphal) response variables of *mtzip14* mutant and segregating wild-type plants either inoculated with the arbuscular mycorrhizal fungus *Rhizophagus irregularis* (pink) or mock-inoculated (blue). The sign and magnitude of the contribution of variables is indicated by the loadings (arrows). The large diamonds signify the mean and 95 % confidence ellipse for each treatment.

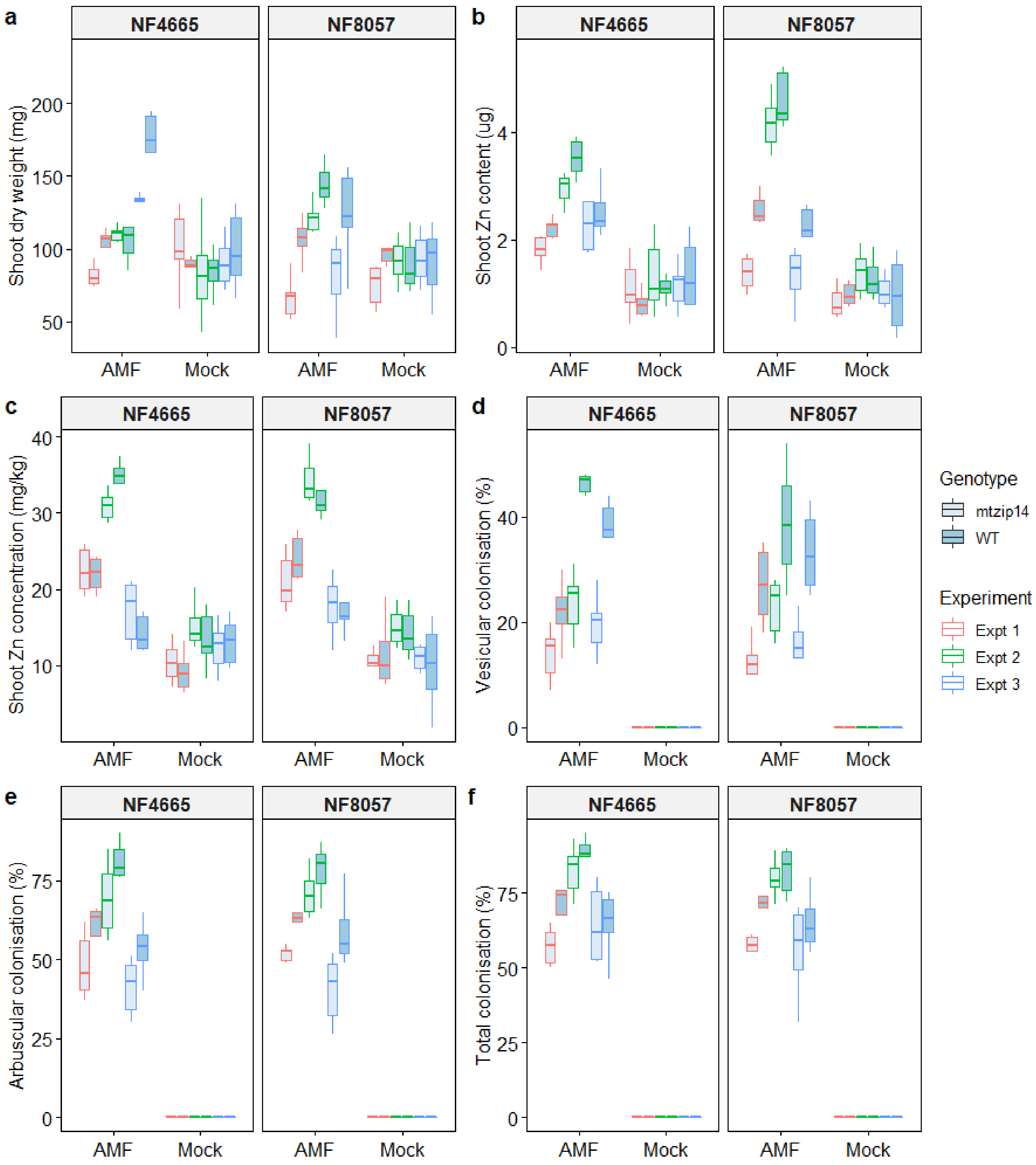


**Figure S8.** Plant biomass (a), zinc nutrition (b-c) and arbuscular mycorrhizal colonisation (d-f) from the *Medicago truncatula mtzip14* and segregating wild-type NF4665 (knock-down) and NF8057 (knock-out) lines from each of the three independent experiments.
